# vardictcpp: A C++ reimplementation of VarDict variant caller

**DOI:** 10.64898/2026.09.23.753771

**Authors:** Joachim Wolff

**Affiliations:** Hannover Medical School, Department of Hematology, Hemostasis, Oncology, and Cell Therapy, Hannover, Germany

**Keywords:** Genomics, Software, Variant Caller

## Abstract

Genomic variant calling is a central computational task in disease research and diagnostics, where somatic mutations such as JAK2 V617F serve as diagnostic markers in myeloid neoplasms. VarDict, originally written in Perl, and its reimplementation VarDictJava are widely used variant callers in the bioinformatics community. The Perl implementation requires run times of up to several hours; VarDictJava reduces this to a few minutes but can consume large amounts of main memory, which limits the number of jobs that can be run in parallel on a given machine. We present vardictcpp, a functionally identical reimplementation of VarDictJava in C++17. vardictcpp is approximately four times faster in single-core execution and 9-11 times faster using eight cores, while requiring up to 17 times less main memory. Variant calls are identical to those of VarDictJava on the evaluated whole-exome samples. The reduced memory footprint allows more concurrent jobs on existing hardware and thereby increases sample throughput, which is particularly relevant for clinical sequencing laboratories with fixed compute resources. vardictcpp is implemented in C++17 and freely available at https://github.com/MHH-Bioinformatics-Hematology/vardictcpp

## 1 Statement of Need

Clinical mutation detection focuses on a defined set of well-characterized variants. Targeted sequencing is normally used for this purpose, although whole-genome sequencing (WGS) and whole-exome sequencing (WES) are also options. In leukemia diagnostics, missense mutations such as JAK2 V617F serve as markers for different disease entities and risk groups [1]. Clinical results must be available within short turnaround times, so any reduction in compute time through a more efficient implementation is beneficial. We use VarDictJava to detect variants at ultra-low variant allele frequencies (VAF), as required for measurable residual disease (MRD) monitoring. Here the VAF threshold of the variant caller must be set to zero in order to detect the faintest traces of malignant cells, which is essential for assessing a potential relapse [2]. Historically, additional parallel capacity could simply be purchased as more hardware, but this is becoming financially increasingly difficult. Driven by demand from artificial intelligence, memory prices have reportedly risen sharply, reaching 448% of the previous year’s level in Germany in the summer of 2026 [3]. Memory efficiency is therefore no longer a secondary concern.

A method to reduce the compute time is to restrict the search to an area of interest. VarDict-Java [4, 5] uses the −*R* parameter to restrict the search to a region. According to the documentation, this parameter is intended for small regions: “The region of interest. In the format of chr:start-end. If chr is not start-end but start (end is omitted), then it is a single position. No BED is needed.” [6]. The documentation does not state whether larger regions are supported. When an entire chromosome is passed to −*R*, however, memory consumption explodes and exceeds that of a genome-wide search with an equivalent BED file, behaviour that is not apparent from the documentation. This usage is not exotic: the Galaxy [7] tool wrapper exposes the region parameter for full chromosomes only [8], so any Galaxy user restricting a run to a chromosome encounters this case. A moderately higher memory usage would be acceptable, however, using this parameter for one full chromosome can lead to a memory consumption of more than 100 GB of RAM.

We therefore reimplemented VarDictJava in C++17 to reduce runtime and, above all, peak memory, so more calling jobs fit on one compute node. This enables a larger number of parallel VarDict calls on the same compute nodes, increasing overall throughput and contributing to faster clinical turnaround times.

## 2 Implementation

vardictcpp is a C++17 reimplementation of VarDictJava [5], developed to provide a functionally equivalent, high-performance alternative for variant calling from next-generation sequencing data. The reimplementation was performed using an agentic AI-assisted translation workflow with Claude Code, in which large language model (LLM) agents iteratively converted Java source modules to idiomatic C++ code, followed by systematic validation against the original VarDictJava output. The agentic AI got as requirements only that the command line interface (CLI) and the input must be identical and the computed output too. How the C++ implementation reaches the identical output was up to the AI. The Java source code can be used as a template, but we did not want an identical source code copy just in another programming language.

vardictcpp depends on `htslib` (v*>*=1.10) for reading and indexing sequence alignment files (BAM/CRAM) and reference FASTA files (via `faidx`), replacing the Java HTSJDK library used in the original implementation. Command-line arguments, default parameter values, and output VCF formatting are designed to exactly mirror those of VarDictJava, allowing vardictcpp to be used as a drop-in replacement in existing pipelines without modification of wrapper scripts. vardictcpp is available on Bioconda (`conda install -c bioconda vardictcpp`) and as a Galaxy tool, or built from source with CMake (≥3.15) and htslib (≥1.10) via `cmake -B build && cmake --build build`.

Each core module of VarDictJava (e.g., pileup construction, soft-clip realignment, structural and complex variant detection) was translated individually and tested against reference outputs generated by running VarDictJava on shared synthetic and real-world BAM datasets. Differences in output between vardictcpp and VarDictJava were logged, triaged, and resolved iteratively until concordant VCF output was achieved across the tested regions. Validation covered every variant class VarDict emits: single-nucleotide variants (SNVs), multi-nucleotide variants (MNPs), insertions, deletions, complex substitutions, and structural variants, under both the default mode (−*f* 0.01) and the ultra-sensitive MRD setting (−*f* 0). Concordance was checked on the complete output row (all 36 columns), so per-strand depths, quality summaries, microsatellite annotation, and genotypes are verified, not only coordinates. The demanding cases were soft-clip realignment, large-indel reconstruction, MNP merging, and multi-variant loci with shared reads. The parity harness is in the repository, and the real BAM inputs and per-sample reports are in the Zenodo archive, so the coverage can be inspected and re-run.

Parallelism follows VarDictJava’s model. The target intervals (a BED file, or a long −*R* region tiled into fixed-size chunks by the same `--chunk` mechanism) are independent work units spread over a pool of worker threads, set with `-th`/`--threads` (default 1). Each interval is processed independently, and the results are re-assembled into VarDict’s deterministic output order before writing. A multi-threaded run is therefore byte-identical to a single-threaded one; thread count changes only runtime and peak memory, not the calls.

Because vardictcpp reproduces rather than replaces VarDict’s calling algorithm, its speed and memory gains come from the execution model rather than the method: running as native compiled code instead of on the JVM removes garbage-collection, JIT-warmup, and boxed-object overhead from the per-read inner loops. The central architectural change is the in-memory representation of per-position evidence, where Java’s graphs of boxed-integer-keyed hash maps referencing heavy-weight variant objects are replaced by primitive-keyed containers holding compact value types, which improves cache locality and lowers peak resident memory by roughly an order of magnitude.

## 3 Results and Discussion

Performance was evaluated on four FLT3-ITD acute myeloid leukaemia (AML) datasets [9] and one MV4-11 AML cell line [10, 11], covering target region counts ranging from 11,000 to 770,000 regions (Table 1). All measurements were performed on an AMD Epyc 9654 (96 cores, 1.1 TB RAM) or an AMD Ryzen 7950X (16 cores, 128 GB RAM) running both Ubuntu 24.04. VarDictJava was benchmarked in its most recent release, version 1.8.3, on a Java Virtual Machine (JVM) version 25. In single-core mode, vardictcpp processed dataset SRR15006376 (100,000 target regions) in 44 seconds using 74 MB of memory, compared with 181 seconds and 1.2 GB for VarDictJava. On SRR15006540 (135,000 target regions), vardictcpp required 61 seconds and 76 MB, against 226 seconds and 1.0 GB for VarDictJava. Across all datasets, the C++ implementation is a factor of 3.7–4.8 faster and requires 8–17 times less memory (Table 2). In multi-core mode, both implementations were restricted to 8 cores. For SRR15006376, vardictcpp required 7.8 seconds and 358 MB, while VarDictJava took 85 seconds and 2.6 GB, the latter still twice as slow as the single-core vardictcpp run. Overall, vardictcpp with 8 cores is a factor of 9.3–11.0 faster than VarDictJava and requires 7–8 times less memory.

**Table 1:** Byte-level parity of vardictcpp against VarDictJava 1.8.3 on five whole-exome sequencing samples (hg19, per-sample covered-target BED, minimum allele frequency 0.01). Sorted output was compared line-for-line against VarDictJava. *Byte-identical* : rows matching the Java reference character-for-character.

| Sample | Target regions | Variant rows | Byte-identical |
| --- | --- | --- | --- |
| SRR15006386 | 11 732 | 4 014 | 4 014 (100%) |
| SRR15006375 | 12 292 | 4 056 | 4 056 (100%) |
| SRR15006376 | 105 949 | 12 739 | 12 739 (100%) |
| SRR15006540 | 135 145 | 16 031 | 16 031 (100%) |
| SRR8657348 | 773 667 | 54 448 | 54 448 (100%) |
| Total |  | 91 288 | 91 288 (100%) |

**Table 2:** Runtime and peak memory of vardictcpp versus VarDictJava 1.8.3 on three whole-exome samples (106–774 k target regions; hg19, minimum allele frequency 0.01) at 1 and 8 threads, CPU-pinned. Median of 5 replicate runs; wall clock and peak resident set size (RSS) from `/usr/bin/time -v`, VarDictJava on JDK 25 with `-Xmx 8g`. *Speedup* = Java */* cpp runtime; *×less* = Java */* cpp peak RSS. Measurement on AMD Ryzen 7950X.

| Sample | Threads | Runtime (s) |  |  | Peak RSS (MB) |  |  |
| --- | --- | --- | --- | --- | --- | --- | --- |
| | | vardictcpp | VarDictJava | Speedup | vardictcpp | VarDictJava | ×less |
| SRR15006376 (105 949) | 1 | 44.5 | 181.1 | 4.1× | 74 | 1274 | 17× |
| | 8 | 7.8 | 85.0 | 11.0× | 358 | 2633 | 7× |
| SRR15006540 (135 145) | 1 | 61.7 | 226.3 | 3.7× | 76 | 1084 | 14× |
| | 8 | 10.4 | 96.9 | 9.3× | 312 | 2569 | 8× |
| SRR8657348 (773 667) | 1 | 98.0 | 473.5 | 4.8× | 122 | 977 | 8× |
| | 8 | 14.8 | 142.4 | 9.6× | 252 | 2084 | 8× |

Memory consumption is by far the highest when the region parameter is set to an entire chromosome. On the MV4-11 AML cell line, restricting the run to the whole of chromosome 1 caused memory usage to explode in both implementations. Instead of 252 MB and 14 seconds with 8 cores (Table 2), vardictcpp required about two minutes (2:03) and 10.6 GB of memory (Table 3). VarDictJava consumed 107 GB of RAM instead of 2 GB and took six minutes rather than two. This corresponds to a 43-fold increase in memory for vardictcpp and a 53-fold increase for VarDictJava. While the increase in the C++ version is substantial, its consumption remains within reasonable bounds, which cannot be said of the Java version, whose 107 GB requirement exceeds the memory available on most compute nodes.

**Table 3:** Peak memory and runtime on a single whole-chromosome region: VarDictJava 1.8.3 (JDK 25, `-Xmx 200g`) versus vardictcpp, both processing chromosome 1 of sample `SRR8657348` (MV4-11 AML cell line, CCLE) as one `-R chr1:1-249250621` work unit, single-threaded and pinned to 8 cores on a 128 GB node. Peak resident set size (RSS) from `/usr/bin/time -v`. At this scale the per-position variation maps span the whole chromosome, so peak memory is driven by region size rather than the minimum allele frequency (−*f* ): `-f 0.01` and `-f 0` blow up equally. VarDictJava does not complete this region at `-Xmx 4g` (out of memory); vardictcpp holds 10.6 GB with no heap tuning.

| Caller | Peak RSS (GB) |  | Runtime (m:ss) |  |
| --- | --- | --- | --- | --- |
|  | -f 0.01 | -f 0 | -f 0.01 | -f 0 |
| VarDictJava 1.8.3 | 107.6 | 107.7 | 6:10 | 6:11 |
| vardictcpp | 10.6 | 10.6 | 2:03 | 2:01 |
| vardictcpp: $\sim 10\times$ less peak memory, $\sim 3\times$ faster. Measurement on AMD Ryzen 7950X. | | | | |

To further demonstrate the performance of the C++ implementation, the speed in single- and multi-core and its memory usage with a VAF of 0.01 was computed for the tools vardictcpp, VarDictJava, FreeBayes [12], LoFreq [13], Mutect2 [14] and octopus [15] on the SRR8657348 dataset with 773,667 target regions. Vardictcpp is by far the fastest tool in single- and multi-core usage. While in single-core it needs 2:21 minutes the next closest one is VarDictJava with around eleven minutes. The other tools with run times of around 5 to 17 minutes are far behind. With 8 cores, vardictcpp completes the analysis in 19 seconds, outperforming all other tools at least by a factor of 2 (FreeBayes, 44 seconds), up to a factor of 59 (octopus, 18:57) see Figure 1 (top). Next, the performance in the somatic mode is measured. Again, vardictcpp is in single- (3:04 minutes) and multi-core (35 seconds) by far the fastest, and needs the least memory. The closest competitor is VarDictJava in single- (9:40 minutes) and multi-core (5:03 minutes), see Figure 1 (bottom). The exact caller commands, target BEDs, and the recorded wall-clock and peak-RSS values behind Figure 1 and Tables 2–3 are in the Zenodo archive (Data Availability), so every comparison can be reproduced.

**Figure 1:**
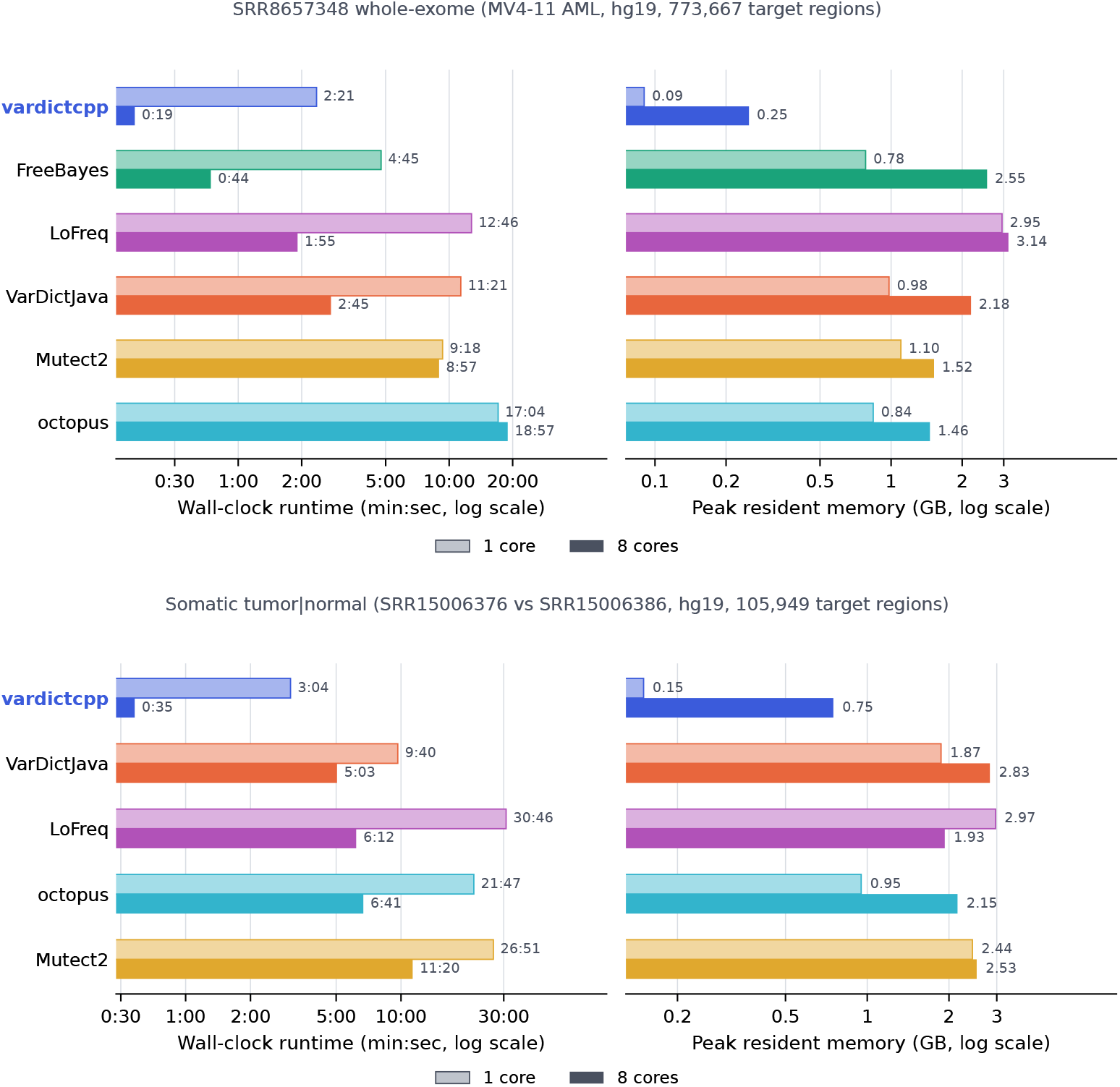
**Top**: Runtime and memory performance for different variant callers for a tumor only sample (SRR8657348) with 773,667 target regions. **Bottom**: Runtime and memory performance in somatic tumor—normal mode using SRR15006376 and SRR15006386. Measurement on AMD Epyc 9654.

Finally, output accuracy was assessed. For all five tested samples in the normal mode, vardictcpp and VarDictJava produced byte-identical output, see Table 1. To further validate, six variant callers were evaluated on chromosome 20 of the Genome in a Bottle (GIAB) germline sample HG002 (Agilent SureSelect v5 exome), scored against the NIST GIAB v4.2.1 truth set with `rtg vcfeval` over the high-confidence, exome-callable regions (depth ≥ 20; 2.31 Mbp; 2 443 truth variants) [16]. While octopus is leading for single nucleotide variants (SNVs) with an F1 score of 0.993, vardictcpp and VarDictJava show identical results and are close to it with 0.985. Considering inserts and deletions (Indels), octopus leads by 0.907, followed by FreeBayes with 0.829 and the two vardict implementations with 0.801, see Figure 2. In somatic mode, we used SRR15006376 as tumor and SRR15006386 as normal input, and the two VarDict implementations return an identical result. While VarDict detects 737 SNVs, Mutect2 125, LoFreq 111 and octopus only 17. However, no orthogonal truth set exists for this tumor—normal pair, so Figure 3 reports inter-caller concordance, not accuracy. The 606 variants called uniquely by VarDict cannot be resolved into true positives and false positives here; they are equally consistent with higher sensitivity at low variant allele frequency and with a higher false-positive rate. The comparison is therefore intended to characterise the calling behaviour of the two VarDict implementations relative to established tools, not to rank callers by accuracy.

**Figure 2:**
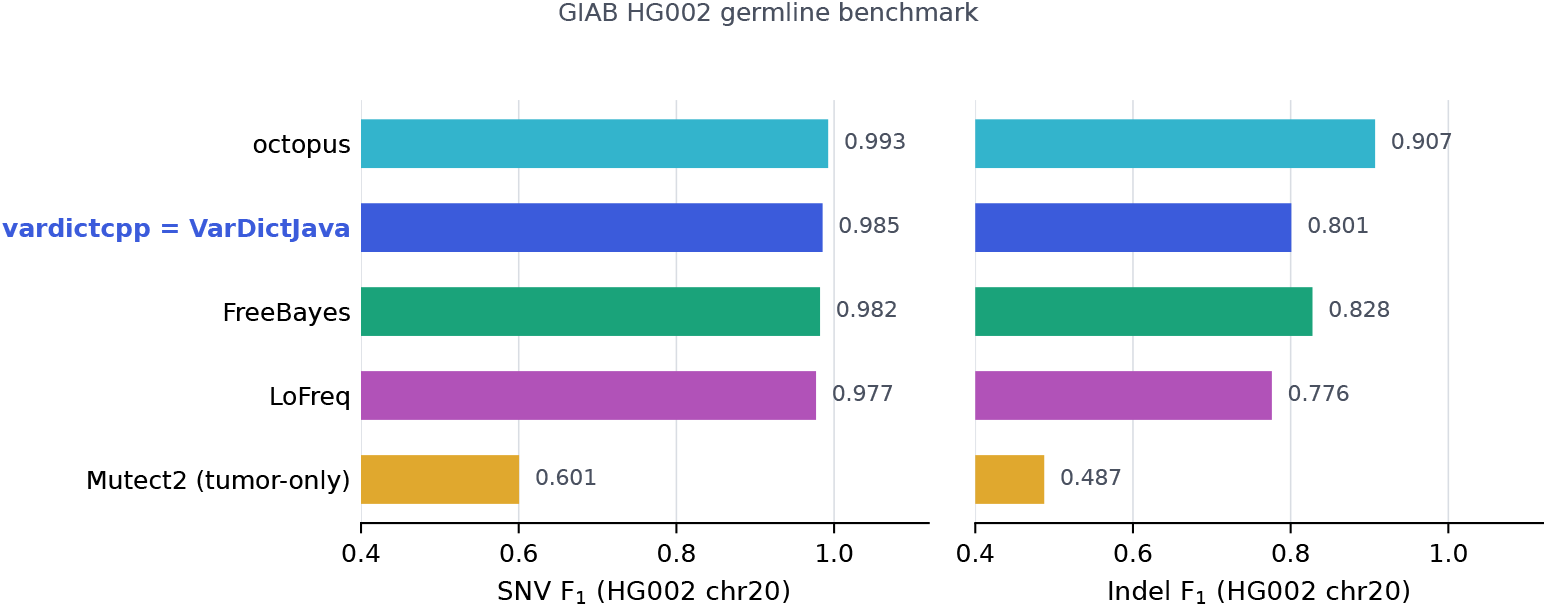
Calling F1 score on chromosome 20 of the Genome in a Bottle (GIAB) germline sample HG002 (see the main text for the truth set and scoring method) [16].

**Figure 3:**
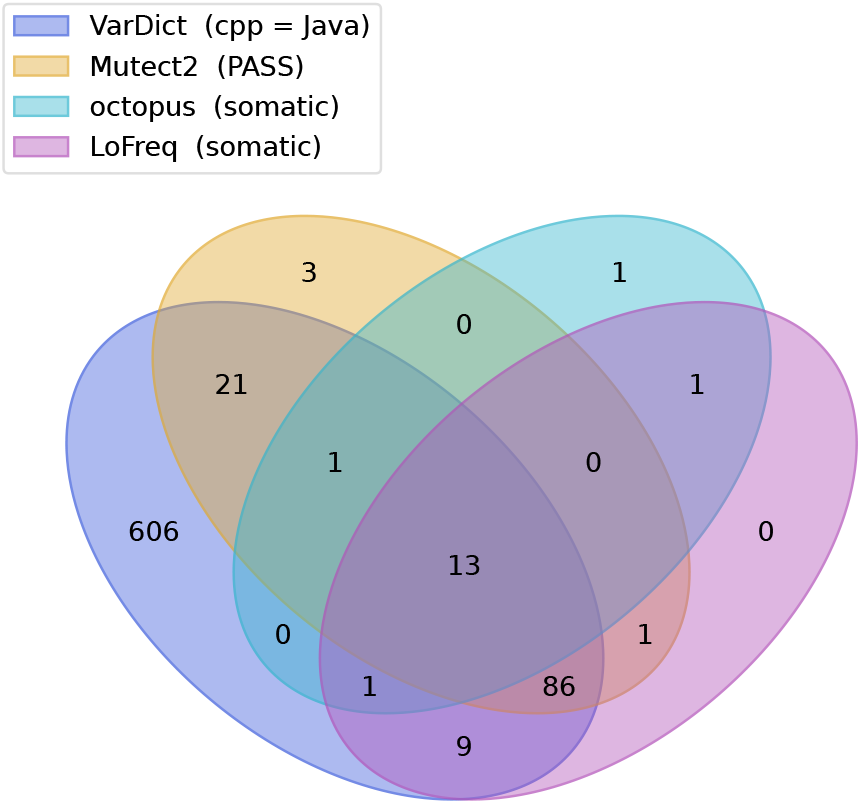
Results of different callers in somatic mode with SRR15006376—SRR15006386 as tumor—normal. Reference genome hg19 and 105,949 target regions. Vardict in Java and C++ give an identical result.

Within this single project, LLM-assisted translation produced a validated, byte-concordant reimplementation in 12 calendar days of part-time work. We do not claim this generalises: the task was unusually well-posed (a fixed CLI, a complete reference implementation, and an automatable byte-level oracle for correctness). Such conditions are common when modernising legacy scientific software, which suggests the approach is worth evaluating more systematically. Recent studies have begun this evaluation: large language models for scientific workflow development [17], and for translating code between programming languages under diverse prompting [18]. For comparison, VarDictJava, itself a port of the original Perl VarDict, and therefore developed from an equivalent validated reference implementation, spanned 143 calendar days from first commit to first release (18 November 2014 to 10 April 2015) [19]. Effort intensity behind the latter figure is unknown, and the two releases may differ in functional scope, so the comparison bounds calendar time rather than developer effort.

## 4 Availability of source code and requirements

- Project name: vardictcpp
- Project home page: https://github.com/MHH-Bioinformatics-Hematology/vardictcpp
- Documentation: https://vardictcpp.readthedocs.io/
- Operating system(s): Linux (x86-64 and ARM), macOS (Apple Silicon)
- Programming language: C++
- Other requirements: CMake *>*= 3.15, htslib
- License: MIT
- RRID: SCR 028800
- Distribution: Bioconda (`conda install -c bioconda vardictcpp`), a Galaxy tool wrapper, or from source with `cmake -B build && cmake --build build`.
- Support: bugs and questions via the GitHub issue tracker at https://github.com/MHH-Bioinformatics-Hematology/vardictcpp/issues.

## 5 Data Availability

Samples SRR15006375, SRR15006376, SRR15006386 and SRR15006540 are adult AML bone marrow samples carrying FLT3-ITD mutations, taken from the study “Frugal alignment-free identification of FLT3-internal tandem duplications with FiLT3r” [9] (BioProject PRJNA742684, Illumina MiSeq). Sample SRR8657348 is the MV4-11 AML cell line from the Cancer Cell Line Encyclopedia (BioProject PRJNA523380) [10, 11].

The intermediate files, such as the BAMs, the reference and truth sets, the caller output files, and the benchmark scripts and results, are available on Zenodo [20]. The source code is available on the project home page.

## 6 Declarations

## 6.1 List of abbreviations

VAF: Variant Allele Frequency
MRD: Measurable Residual Disease
AML: Acute Myeloid Leukemia
ITD: Internal Tandem Duplication
WGS: Whole Genome Sequencing
WES: Whole Exome Sequencing
RSS: Resident Set Size
JVM: Java Virtual Machine
LLM: Large Language Model
VCF: Variant Call Format
CLI: command line interface

## 6.2 Ethical Approval

All data analysed are publicly available under BioProjects PRJNA742684 and PRJNA523380; no new human data were generated. No ethical approval was required.

## 6.3 Competing Interests

The author declares that they have no competing interests.

## 6.4 Author’s Contributions

J.W. implemented the project, supervised the agentic AI and wrote the paper.

## 6.5 Use of generative AI in manuscript, figure generation and software

Anthropic Claude Code has been used to generate the source code of vardictcpp. Models used were Claude Opus 4.8. Moreover, Claude Opus 4.8 has been used in assistance to write the text, and to correct the grammar. Please refer also to the commit history on GitHub. Time of usage: July 22 - August 07, 2026 [21].

## Notes

https://doi.org/10.5281/zenodo.21836659

### Competing Interest Statement

The authors have declared no competing interest.

